# The cost of sympatry: spatio-temporal patterns in leopard dietary and physiological responses to tiger competition in Rajaji Tiger Reserve, India

**DOI:** 10.1101/2022.08.03.502614

**Authors:** Shiv Kumari Patel, Sourabh Ruhela, Suvankar Biswas, Supriya Bhatt, Bivash Pandav, Samrat Mondol

**Affiliations:** Wildlife Institute of India, Chandrabani, Dehradun, Uttarakhand 248001

**Keywords:** Inter-species competition, Dominance hierarchy, Habitat recovery, Physiological cost, Stress and nutrition hormones, Carnivore conservation

## Abstract

1. Apex predators have critical roles in maintaining the structure of ecosystem functioning by controlling intraguild subordinate populations. Such dominant-subordinate interactions involve agonistic interactions including direct (death/displacement) or indirect (physiological and/or health implications) impacts on the subordinates. As these indirect effects are often mediated through physiological processes, it is important to quantify such responses for better understanding of population parameters.
2. We used a well-known sympatric large carnivore intraguild system involving tiger (*Panthera tigris*) and leopard (*Panthera pardus*) to understand the dietary and physiological responses under a spatio-temporal gradient of tiger competition pressures in Rajaji Tiger Reserve (RTR), a major source tiger population of the western Terai-Arc Landscape, India between 2015-2020. The park provided a natural experimental set-up for tiger competition with the eastern part (ERTR) having high tiger density and the western (WRTR) part with functionally no competition from tigers.
3. We conducted systematic faecal sampling in the winters of 2015 and 2020 from ERTR and WRTR to assess diet and physiological measures. Analyses of leopard-confirmed faeces suggest a dietary-niche separation as a consequence of tiger competition. In 2020, we found increased occurrence of large-bodied prey species without tiger competition in WRTR. Physiological measures followed the dietary responses where leopards with large-sized prey in diet showed higher fT3M and lower fGCM measures in WRTR. In contrast, ERTR leopards showed lower levels of fT3M as well as fGCM in 2020, possibly due to intense competition from tigers. Overall, these pattens strongly indicate a physiological cost of sympatry where competition with dominant tigers resulted in elevated nutritional stress.
4. The combination of the natural habitat providing unique experimental setup, spatio-temporal sampling strategy and multidisciplinary approaches provide critical conservation perspectives for leopards, particularly in the context of recent increase in tiger numbers across India. We recommend expansion of leopard monitoring and population estimation efforts to buffers, developing appropriate plans for human-leopard conflict mitigation and intensive efforts to understand leopard population dynamics patterns to ensure their persistence during the ongoing Anthropocene.

## 2. Introduction

Apex predators are critical in maintaining the structure and control of the local ecosystem functioning through prey-predator dynamics (Ritchie & Johnsan 2009; Ritchie et al., 2012; Gaynor et al., 2019; Ripple et al., 2014) and their limiting effects on subordinates (Ritchie & Johnsan 2009; Feit et al., 2019; Suraci et al., 2016; Ripple et al., 2014). Intensity of such limiting effects within a large carnivore guild is more pronounced for species competing for similar resources (Palomares & Caro 1999; Donadio & Buskirk 2006; Ritchie & Johnson 2009). Dominant species within the guild exercise interference competition through aggression (Merkle et al., 2009; Linnell & strand 2000), harassment (Merkle et al., 2009), kleptoparasitism (Périquet et al., 2015; Merkle et al., 2009), delaying intraspecific communication (Cornhill & Kerley 2020) and direct killing (Palomares and Caro 1999; Ritchie & Johnsan 2009; Donadio and Buskirk 2006; Laurenson 1994; Merkle et al., 2009). Subordinate members, on the other hand employ various tactics (for example, spatio-temporal and dietary separation) to minimize interference and maximize resource acquisition (Hayward et al., 2009, Vanak et al., 2013) to achieve a balance for successful co-existence. Local ecological factors are known to drive such behavioral strategies, which has been an area of extensive research interest in various carnivore guilds globally (López-Bao et al., 2016; Périquet et al., 2014; Vanak et al., 2013; Karanth & Sunquist 1995; 2000; Steinmetz et al., 2013; Broekhuis et al., 2013; Carlsson et al., 2010).

Dominant-subordinate agonistic interactions exerts two different kinds of negative impacts on the subordinate species: (a) direct impacts such as getting killed or displacement from the best-quality habitats (Newsome et al., 2017; Ramesh et al., 2017; Merkle et al., 2009; Mitchell & Banks 2005) and (b) indirect effects from increased pressures from competitions, inadequate food resources and resulting energy deficits (Sheriff et al., 2020; Creel et al., 2008, 2017; Suraci et al., 2016) affecting survival, growth, body condition, reproduction and parental provisioning (LaManna and Martin 2016; Parker et al., 2009; Creel et al., 2007; 2008; 2009). As many of these indirect effects are mediated through physiological processes (Macleod et al., 2018; Clinchy et al., 2013), quantification of the physiological responses is essential to understand changes in various population parameters of the subordinate species (Creel et al., 2009; Gaynor et al., 2019). Recent advances in physiological measurements of environmental stressors, particularly in combination with non-invasive sampling approaches have made it easier to link the environmental effects with their respective physiological responses (Ames et al., 2020; Palme 2019; Sopinka et al., 2015; Dantzer et al., 2014). For example, a number of inter-species (predator-prey-see Ylönen et al., 2006; Creel et al., 2009, dominant-subordinate dynamics-see Van Meter et al., 2009 etc.) and intra-species (social hierarchy-see Sapolsky 1983; Armitage 1991; Creel et al., 1996; Creel 2001; Van Meter et al., 2009, competition-see Armitage 1991; Creel 2001) interactions have been addressed using glucocorticoid (GC) measures, demonstrating its use. Further, recent addition of thyroid hormone (T3, in particular) (Behringer et al., 2018; Wasser et al., 2010; Flier et al., 2000; Eales, 1988) measure is allowing us to separate the impacts of dietary resource availability from overall stress measures (through GC) as shown in marine (McCormley et al., 2018; Wasser et al., 2017; Ayres et al., 2012) and terrestrial mammals (Szott et al., 2020; Dias et al., 2017; Joly et al., 2015; Vynne et al., 2014; Wasser et al., 2011), including large carnivores (Patel et al., 2021; Vynne et al., 2014).

The sympatric tiger (*Panthera tigris*) and leopard (*Panthera pardus*) are one of the most well-studied model systems to understand the dominant-subordinate intraguild competition (Seidensticker 1976; Mcdougal 1988; Mondal et al., 2012; Steinmetz et al., 2013; Carter et al., 2015; Pokheral & Wegge 2019; Kafley et al., 2019; Kumar et al., 2019; Thapa et al., 2021). Leopards, when co-existing with tigers, are often dominated by their larger counterpart in terms of resources (both space and food) (Seidensticker 1976). Large number of studies have focused on exploring different strategies adopted by leopards such as spatio-temporal (Thapa et al., 2021; Kafley et al., 2019; Kumar et al., 2019; Pokheral & Wegge 2019; Carter et al., 2015) and dietary niche segregation (Pokheral & Wegge 2019; Harihar et al., 2011; Andheria et al., 2007; Karanth & Sunquist 1995) for successful co-existence with tigers, but the physiological consequences of such interactions have received less attention. Here we address leopard physiological and dietary responses in the context of competition with tigers within Rajaji Tiger Reserve (RTR), western Terai-Arc landscape, India. RTR is a major source tiger population (estimated density of 8 ±1.4/100 km^2^ in 2018) and retains one of the highest density of leopards (16.90 ±1.44/100km^2^) in the landscape (Jhala et al., 2021). The park is physically separated by the river Ganges in two parts: eastern and western RTR (henceforth ERTR and WRTR) that are structurally connected by a narrow corridor, heavily affected by anthropogenic activities (Johnsingh et al., 1990; Harihar et al., 2018; Biswas et al. 2022a) (Figure 1). Both sites are similar in terms of wild prey densities and vegetation structure (Harihar et al., 2009b) but differ in the extent of tiger competition intensity. Almost the entire tiger population of RTR is found in the ERTR whereas leopard, in the absence of tiger, is functionally the apex predator in the WRTR. This unique situation provides an ideal, natural system to assess the physiological impacts of inter-species competition in a control-test framework (WRTR is a control site with no inter-species competition when compared with ERTR). We used leopard faecal hormone metabolite measurements (fGCM and fT3M) in 2015 and 2020 to address spatio-temporal differences in physiological and dietary responses against a tiger competition gradient. More specifically, we asked the following questions: (i) how leopard dietary profiles, fGCM and fT3M measures vary with changing tiger competition intensities over space and time, and (ii) how local ecological factors (habitat productivity and prey body size) explain such differences in leopard physiology. We believe that results of this study have larger implications in understanding the physiological costs for subordinate carnivores co-existing within a guild and their long-term conservation.

**Figure 1.**
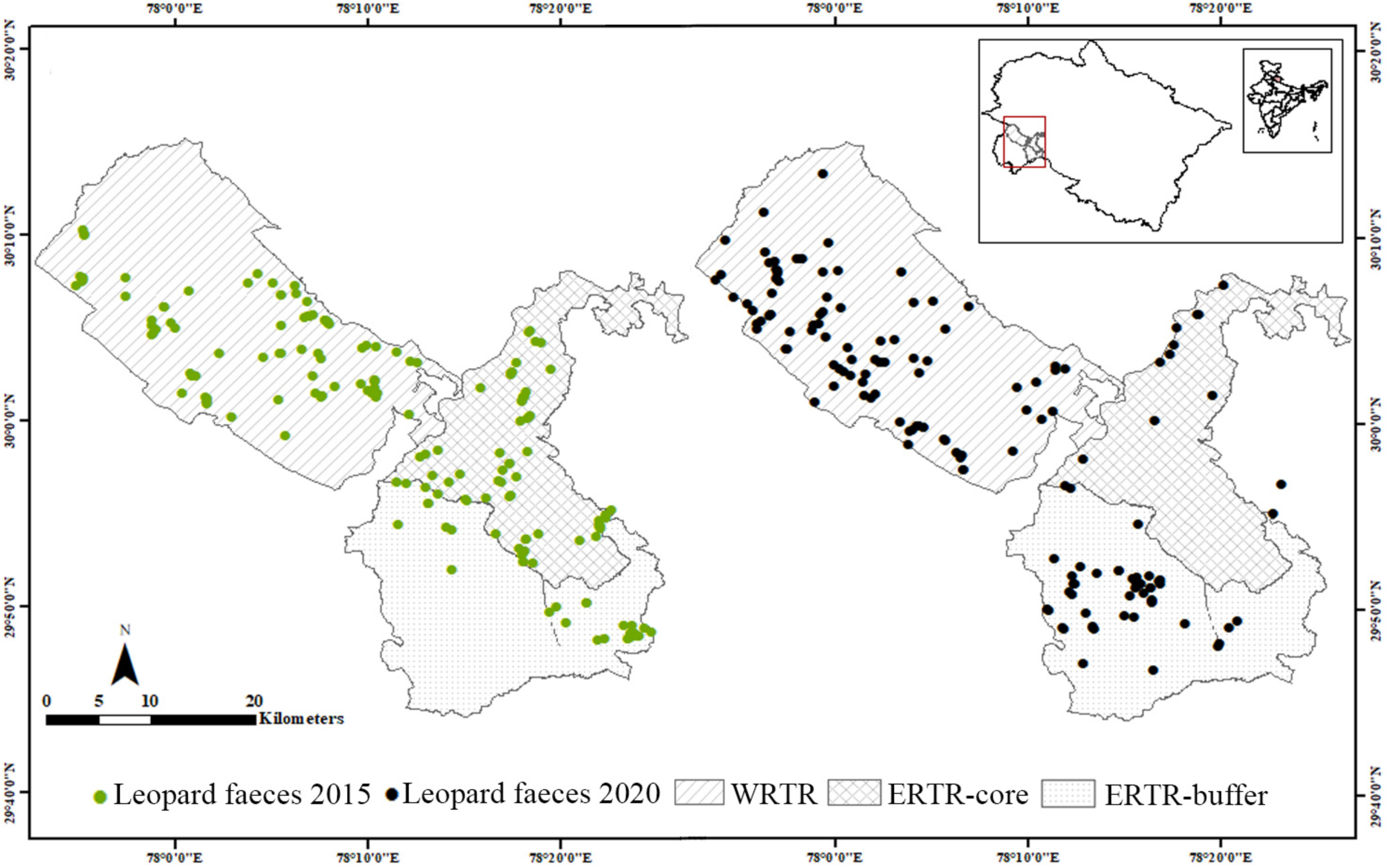
Spatio-temporal locations of field-collected leopard faecal samples from Rajaji Tiger Reserve (RTR) used in this study. The core and buffer zones are differentially marked. The left pane represents samples collected in 2015, whereas the right pane shows 2020 samples.

## 3. Material and methods

### 3.1 Research permissions and ethical considerations

All required permissions for field surveys and faecal sampling were provided by Forest Departments of Uttarakhand ((Permit no: 90/ 5–6) and (Permit no: 3707/ 5–6)). Due to the non-invasive nature of this study, no further ethical clearance was required.

### 3.2 Study area

This study was conducted in Rajaji Tiger Reserve (RTR) (Figure 1), the westernmost part of Rajaji-Corbett Tiger Conservation Unit (RCTCU, Johnsingh & Negi 2003), in the Indian part of the Terai-Arc landscape (TAL). Located at base of Himalayan foothills and starting of Indo-Gangetic plains, RTR has an undulating topography with a mosaic of woodlands and grasslands, drained by multiple rivers and streams. The forest type is broadly classified as northern Indian moist deciduous, dominated by Sal (*Shorea robusta*) (Champion and Seth, 2005). The park is naturally separated by the river Ganges in two parts, connected by a narrow corridor called as Chilla-Motichur corridor (3 km length and 1 km width) (Johnsingh et al., 1990; Harihar et al., 2018). The eastern part (covering 579 km^2^ area of core and buffer zones), situated in the east bank of Ganges maintains structural as well as functional connectivity with major tiger populations (such as Corbett Tiger Reserve) of the landscape (Harihar et al., 2020; Biswas et al. 2022a). However, the western part (571 km^2^ area on the west bank of Ganges) has become isolated over years from the ERTR. Historically, the entire study area (RTR) was inhabited by large numbers of agro-pastoralist community settlements (Toungya and Gujjars, respectively) which were primarily dependent on forest resources (Berkmuller et al., 1987), leading to forest degradation. In 1983, Rajaji was declared as a national park and significant efforts towards tiger and prey population recovery were undertaken. As part of this, a plan for relocation and rehabilitation of local communities was prepared to create inviolate space for wildlife (Roy 2003), and accordingly all settlements from the core areas of ERTR were relocated by 2003 (Harihar et al., 2009b). Subsequently, the conditions of the natural habitat improved and the densities of the wild prey species increased and facilitated population recovery of wild tigers in ERTR (from 2.08 in 2004-5 to 7.05 individuals/100km^2^ in 2016-17, Harihar et al., 2020). However, in the WRTR, the human rehabilitation was completed in periodical manner where by 2005 some ranges were made inviolate (Harihar et al., 2009b) and others were completed by 2016. As WRTR remained functionally disconnected from eastern counterpart during this period, the tiger population reduced drastically (from 5-10 individuals in 2000 to two females in 2006, Johnsingh et al., 2006, Jhala et al., 2020) despite comparable prey density (41.22±6.65 individuals/km^2^ in WRTR and 39.23±4.76 individuals/km^2^ in ERTR, Harihar et al., 2020). The leopard population, on the other hand, showed a reverse trend where ERTR recorded a decrease in their density (9.76 to 2.07 individuals/100km^2^ between 2004 to 2008, respectively, Harihar et al., 2011) (possibly due to inter-species competition) and WRTR showed an increasing trend in leopard density (Jhala et al., 2021) (due to less competition from tigers). Such contrasting population patterns under a scenario of inter-species competition provided an ideal ‘natural experimental setup’ to understand leopard physiological responses under low and high tiger density in adjacent and similar habitats. Further, we conducted this study over a five-year temporal framework (2015 and 2020) during which the ERTR has seen a significant increase in tiger population (2.90±0.87 to 8±1.4/100 km^2^, Jhala et al., 2015 and 2020), therefore providing an opportunity to test the effects of competition against a tiger density gradient (high tiger density in 2020 than 2015 within ERTR).

### 3.3 Study design

The unique situation of naturally high (ERTR) vs. low (WRTR) tiger density and reverse patterns of leopard density in adjacent and similar habitats allowed us to ask some important questions regarding various effects of inter-species competition. From a spatial difference perspective we hypothesize that: (1) there would be a dietary niche separation between ERTR and WRTR, where the eastern leopard population would show higher frequency of medium and small sized prey in their diet than the western population (where they don’t face competition from tigers); (2) corresponding high fGCM in ERTR (due to inter-species competition) than western population; and (3) lower fT3M in ERTR leopards (due to possible dietary niche separation) than their western counterpart. In a temporal data perspective, we expect that (4) there would be no significant difference in prey relative frequency of occurrence (RFO) in diet, as well as in fGCM and fT3M measures in WRTR; whereas (5) significant difference in prey RFO in diet, fGCM and fT3M is expected in ERTR resulting from increased competition from tigers between 2015 and 2020 (tiger density of 2.90±0.87/100 km^2^ in 2015 and 8±1.4/100 km^2^ in 2020, Jhala et al., 2020). Apart from the inter-species competition we also expected habitat productivity-related differences in leopard dietary and physiological responses. We expected better habitat to be associated with higher frequency of large sized prey in diet, lower fGCM and higher fT3M levels.

### 3.4 Faecal sampling, species confirmation and prey identification

Due to the spatio-temporal nature of the study, it was critical to establish a standard sampling framework for faecal collection from the entire study area. Some of the major concerns were identification of sampling trails across RTR, seasonal effects, uniform field efforts, constant sampling team etc. As RTR has been a long-term study site for photographic monitoring of tigers (Harihar et al., 2009a; 2009b; 2012; 2020) and earlier genetic studies have used already identified forest trails and tracks (Biswas et al., 2022a), the information was used for faeces collection in this study during both sampling periods (2015 and 2020). Two experienced research and tracking team (each consisting of 3-4 members) surveyed all identified forest trails during 2015 and 2020 and collected fresh faecal samples of large carnivores. Both sampling (in 2015 and 2020) were conducted in winter (December-January) to counter seasonal variations and make use of better environmental conditions (mean temperature of 4-6°C) that maintain relatively long-term sample freshness. In field, sample freshness was determined based on intactness, minimal insect activity and strong odor (Vynne et al., 2014). All fresh faecal samples were collected in wax paper with location details and stored in zip-lock bags (Biswas et al 2019) before transporting them to the laboratory, where they were stored −20 °C until laboratory analysis.

In the laboratory, the samples were genetically ascertained using leopard-specific molecular markers (Mondol et al., 2014) to ensure only confirmed leopard samples were used in downstream diet and physiological analyses. In brief, DNA extraction was performed using a modified Qiagen DNA extraction protocol (Biswas et al. 2019) for all samples and leopard-specific mitochondrial DNA markers (TigParND4-F and ParND4-R, Mondol et al., 2014) were used to ascertain leopard faeces. Confirmed leopard samples were further dried at 60 °C for 72 hours in an oven (#Unilab-112HO, Haryana, India) to control for moisture (Wasser et al., 1993). The undigested parts (prey hair, broken bones, hoof etc.) were separated by sieving the dried samples through sterile 0.5 mm stainless steel mesh and the faecal powders were collected and stored in −20 °C. The primary guard hairs (20-30 hairs/sample) were used to prepare permanent slides and were examined for medulla structures (Mukherjee et al. 1994) using available references (Bahuguna et al. 2010, Biswas et al. 2022b) to identify leopard prey species. Sample size estimation for diet analyses was conducted through a sample rarefaction curve (Magurran 2004), where the species diversity in leopard diet was estimated using Shannon diversity index (Magurran 2004) with EstimateS (Colwell 2006).

### 3.5 Habitat productivity assessment

For leopards or for large carnivores in general a good quality habitat is one with good prey availability (Carbone & Gittleman 2002). Prey availability is associated with forage availability that is often quantified in terms of vegetation cover or green cover (Pettorelli et al., 2005a, b, 2011). We used vegetation cover as a proxy of habitat productivity (Pettorelli et al., 2011), that would facilitate higher prey base for leopards and quantified it by extracting Normalized Difference Vegetation Index (NDVI) values. We used 16-day composite NDVI values recorded by NASA’s MODIS (Moderate Resolution Imaging Spectroradiometer, MOD13Q1 version 6.1 at 250m resolution), downloaded for RTR (for the month of December, corresponding to winter sampling season) for year 2015 and 2020. The analyses were conducted at two scales: (a) For overall habitat productivity assessment, we divided study area into three zones: WRTR, ERTR-core and ERTR-buffer (see figure 1). Each zone was further divided into 3km X 3km grids (9 km^2^ area, approximate leopard home range see Seidensticker 1990); and (b) For sample-based assessment, we used leopard faecal sample location as center and drew buffers of 2 km radius (12 km^2^ area) around each faecal sample. We extracted mean NDVI values for each grid and buffer using MODIS raster images of year 2015 and 2020, where extraction was done using zonal statistics tool (as table for grids and table 2 for buffers) in ArcMap 10.5 (ESRI 2016).

### 3.6 Hormone metabolite extraction and assays

Recent study on wild tigers in the same landscape showed highly variable contents of inorganic matter (IOM) in the faeces that negatively impacted fGCM and fT3M measures (Patel et al., 2021). As leopards share the same space, environmental conditions and prey base in RTR, the field-collected samples were processed for percent IOM measures using the same approach described in Patel et al (2021). In brief, 0.1g of faecal powder was ashed in a muffle furnace (#NSW-101, NSW, New-Delhi, India) at 550^◦^C for 2 h, reweighed and the amount of IOM was calculated. As suggested in the earlier study (Patel et al., 2021), leopard samples with <80% IOM content were used for hormone assays.

For hormone metabolite extractions, each dried faecal powder was thoroughly mixed and 0.1 grams of powder was weighed. The extraction procedure involved pulse-vortexing the weighed amount of faecal powder in 15 ml of 70% ethanol for 30 minutes, followed by centrifugation at 2200 rpm for 20 min (Mondol et al., 2020; Wasser et al., 2010). The hormone extracts were collected in 2 ml cryochill vials (1:15 dilution) and stored at −20 °C in freezer until assays. Leopard fGCM and fT3M were measured using Corticosterone (#K014, Arbor Assays, MI, USA) and Triiodothyronine (T3) (#K056, Arbor Assays, MI, USA) EIA kits. These kits were earlier successfully validated in wild tigers (from TAL) and captive lions (Patel et al., 2021, Goswami et al., 2021) and thus were considered suitable for this study. Further, parallelism and accuracy tests were conducted for leopard faecal extracts in the laboratory. Serial dilutions of faecal extracts paralleled well with standard curves of fGCM (Supplementary figure 1a) as well as fT3M (Supplementary figure 1c). F ratio test showed no differences between slopes of standard and pooled extract curves for fGCM (F(1,10) = 1.89, P = 0.2) and fT3M (F(1,11) = 1.34, P = 0.27). Accuracy tests using regression analysis produced slopes of 1.09 and 1.02 at working dilution of 1:120 and 1:7.5 for fGCM and fT3M (Supplementary figure 1b and 1d). Intra-assay coefficient of variation (CV) was 7.15 and 8.36, whereas inter-assay CV was 10.35 and 7.86 for fGCM and fT3M, respectively. During assays, hormone extracts were dried and reconstituted in assay buffers at required dilution (1:120 for fGCM and 1:7.5 for fT3M). Samples were assayed in duplicate using kit protocols and optical density (at 450 nm) was measured with ELISA plate reader (#GMB-580, Genetix Biotech Asia, New Delhi, India). Cross-reactivities of respective antibodies are presented in supplementary table 1.

### 3.7 Statistical analysis

The analytical framework was established based on the hypothesis proposed in this study, where comparisons were made at two scales: 1) at spatial level, prey and hormone metabolite (fGCM and fT3M) data between ERTR vs. WRTR (individually the 2015 and 2020 data); and 2) at temporal scale where comparisons were made with each part of RTR (2015 vs 2020 for ERTR and WRTR, respectively). While reporting the methods and results following terms have been used to describe the sampled groups: ERTR in 2015- ERTR_2015_, ERTR in 2020- ERTR_2020_, WRTR in 2015- WRTR_2015_ and WRTR in 2020- WRTR_2020_.

To ascertain leopard food habit, data on relative frequencies of occurrences (RFO) for each prey species was calculated using formula i/j*100, where ‘i’ represents the frequency of number of samples in which a specific prey occurs and ‘j’ represents the total frequency count of all prey species (Mukherjee et al. 1994, Kruuk 1989). Further, Relative prey biomass consumed was calculated using formula D= (A*Y)/∑(A*Y)*100 where, ‘A’ represents the RFO of each prey species and ‘Y’ represents weight of consumed prey in each faeces. ‘Y’ is calculated using Ackerman’s equation: Y=1.980 + 0.035X, where X=mean body weight of a particular prey species (Ackerman et al. 1984, Karanth & Sunquist 1995). The mean body weight of prey was taken from Harihar et al. (2011), Rathore (2015) and Upadhyaya et al. (2018) (Table 1). Two-way ANOVA was used to test any significant differences in prey RFO and biomass among sampled groups (Spatial scale: ERTR_2015_ vs. WRTR_2015_, ERTR_2020_ vs. WRTR_2020_; temporal scale: ERTR_2015_ vs. ERTR_2020_, WRTR_2015_ vs. WRTR_2020_). Additionally, all the prey species data was categorized into three major classes: (a) large (≥60 kgs), (b) medium (between 16-60 kgs); and (c) small (≤15 kg) and absolute frequency of occurrence (AFO) was calculated for these classes using formula s*_k_*\*100/n, where ‘s*_k_*’ is the number of samples containing class ‘*k*’ and ‘n’ is total number of faeces analyzed (Harihar et al., 2011). Any difference in AFO percentage in sampled groups were tested using chi-square analyses, followed by f-test for pair wise comparison at spatial and temporal scales (mentioned above). For overall assessment of the habitat productivity, we compared mean NDVI values of three zones (WRTR, ERTR-core and ERTR-buffer) spatially using one-way ANOVA (with subsequent post hoc Tukey’s HSD test) and temporally (between 2015 and 2020) using paired t-test. All analyses were conducted using SPSS version 20 (IBM, 2011).

**Table 1.** Details of the various leopard diet parameters for all nine prey species identified in this study. Results are presented for percentage of relative frequency of occurrence (% RFO), and relative biomass of the consumed prey species in ERTR and WRTR for both 2015 and 2020, respectively.

| Prey species | Mean body weight of prey (in kg) | ERTR<sub>2015</sub> (n=86) |  | WRTR<sub>2015</sub> (n=76) |  | ERTR<sub>2020</sub> (n=53) |  | WRTR<sub>2020</sub> (n=89) |  |
| --- | --- | --- | --- | --- | --- | --- | --- | --- | --- |
|  |  | RFO (%) | Relative biomass (%) | RFO (%) | Relative biomass (%) | RFO (%) | Relative biomass (%) | RFO (%) | Relative biomass (%) |
| Livestock | 250 | 5.81 | 12.50 | 0.00 | 0 | 1.89 | 4.32 | 0.00 | 0 |
| Sambar | 185 | 19.19 | 32.51 | 20.39 | 37.54 | 19.81 | 35.75 | 31.46 | 48.39 |
| Nilgai | 169 | 6.98 | 11.04 | 4.61 | 7.91 | 5.66 | 9.54 | 10.67 | 15.33 |
| Chital | 50 | 38.72 | 28.95 | 48.68 | 39.53 | 47.17 | 37.55 | 43.26 | 29.35 |
| Wild pig | 35 | 4.65 | 2.99 | 5.26 | 3.67 | 5.66 | 3.87 | 3.93 | 2.29 |
| Hog deer | 33 | 7.56 | 4.75 | 7.24 | 4.94 | 0.00 | 0 | 2.81 | 1.60 |
| Langur | 10 | 0.58 | 0.27 | 0.66 | 0.33 | 0.00 | 0 | 0.00 | 0 |
| Hare | 4 | 13.72 | 5.83 | 13.16 | 6.07 | 19.81 | 8.96 | 7.30 | 2.82 |
| Indian peafowl | 5 | 2.67 | 1.16 | 0.00 | 0 | 0.00 | 0 | 0.56 | 0.22 |

During hormone data analyses, the leopard fGCM and fT3M data (raw as well as log transformed) were assessed for normality using diagnostic plots (density plots) and Shapiro-wilk test. Generalized linear models (GLMs) with log link and gamma distribution errors were used to explain the variation in fGCM and fT3M data. To assess any possible changes in fGCM and fT3M levels across spatial (ERTR vs WRTR) and temporal (2015 vs 2020) scales, an interaction term ‘Area*Year’ (as the tiger density in ERTR was lower in 2015 than in 2020, that may impact fGCM and fT3M levels) and the prey size class (large, medium and small, as prey size may impact fGCM and fT3M levels) were used as explanatory variables. Likelihood ratio test (LRT) was used to determine if the explanatory variables explain the data independently or in combination. Finally, post-hoc Tukey’s HSD test was employed to assess any pair-wise differences in fGCM and fT3M levels for all sampled groups (mentioned above) and prey size classes. To evaluate the relationship of fGCM and fT3M with habitat productivity (NDVI values derived from sample buffers), we used two separate linear models (function ‘lm’). All analyses were conducted in R v4.1.1 (R Core Team, 2021) with the following packages: ‘ggpubr’ (Kassambara 2020), and ‘multcomp’ (Hothorn et al., 2022).

## 4. Results

During the study period, a total of 564 large carnivore faeces was collected (n=276: ERTR-172 and WRTR-104 samples in 2015 and n=288: ERTR-178 and WRTR-110 samples in 2020) from the entire study area. After species confirmation, 324 leopard faecal samples were further processed for dietary and hormone analyses. The distribution of these samples was as followed: ERTR_2015_- 92, WRTR_2015_- 81, ERTR_2020_- 60 and WRTR_2020_- 91. However, prey species could be identified from 304 samples (93.82% success rate, ERTR_2015_- 86, WRTR_2015_- 76, ERTR_2020_- 53 and WRTR_2020_- 89, respectively; Table 1). The remaining samples (n=20, 6.18%) contained damaged hairs for accurate species identification and were excluded from further dietary analyses. For hormone analyses, samples with >80% IOM were discarded (n=121) and finally 203 faecal hormone extracts (ERTR_2015_- 56, WRTR_2015_- 49, ERTR_2020_- 42 and WRTR_2020_- 56) were used in physiology analyses.

### Food habits of leopard

Overall, a total of nine prey species (large-bodied-Sambar, Nilgai and Livestock; medium-bodied-Chital, Wild pig and Hog deer and small-bodied- Langur, Hare and Peafowl) were detected. The large and medium-bodied prey species contributed 85.38% (RFO) of leopard diet whereas small prey species comprised only 14.62% (RFO). RFO of Chital (44.49%) and Sambar (22.71%) were highest followed by others (Table 1). However, biomass of sambar was highest (38.55%), closely followed by chital (33.85%) (Table 1). Majority of the samples (N=273, 89.8%) contained single prey species. All prey species except livestock (identified only in the ERTR) was found across all sampled groups. The rarefaction curve saturated beyond 40 samples within each group and no new prey species was identified further (supplementary figure 2).

The two-way ANOVA analyses with all leopard prey species among the sampled groups (Spatial: ERTR_2015_ vs. WRTR_2015_, ERTR_2020_ vs. WRTR_2020_; Temporal: ERTR_2015_ vs. ERTR_2020_, WRTR_2015_ vs. WRTR_2020_) showed no significant differences in both RFO and biomass. However, overall comparisons within habitats across all compared groups indicated significant differences in both RFO and biomass (Supplementary Table 5). Comparative analyses (Chi-square test with prey body-size groups) revealed large-sized prey frequencies differed significantly among sampled groups (χ^2^=8.62, P=0.035). F-test showed that at spatial scale, there were no significant differences in frequencies of different prey classes between ERTR_2015_ and WRTR_2015_ (p = 0.445, p = 0.476 and p = 1.00 for large, medium and small prey classes, respectively) (Table 2). However, compared to ERTR_2020,_ WRTR_2020_ showed significantly higher large-bodied (p = 0.018) and lower small-bodied prey species (p = 0.019). Temporally, the ERTR_2015_ and ERTR_2020_ showed no difference in prey class occurrences (p = 0.445 (large), p = 1.00 (medium) and p = 0.335 (small), whereas WRTR_2020_ showed a significant increase in large prey occurrences (p = 0.018) with no differences in medium and smaller-bodied prey classes (p = 0.156 and p = 0.238, respectively) compared to WRTR_2015_.

**Table 2.** Spatio-temporal comparison of differences in absolute frequency of prey occurrence in leopard diet at Rajaji Tiger Reserve, India. The comparisons were calculated using Fisher’s Exact test (2×2).

| Prey category | Detection | Spatial comparison |  |  |  |  |  | Temporal comparison |  |  |  |  |  |
| --- | --- | --- | --- | --- | --- | --- | --- | --- | --- | --- | --- | --- | --- |
|  |  | ERTR<sub>2015</sub> | WRTR<sub>2015</sub> | P | ERTR<sub>2020</sub> | WRTR<sub>2020</sub> | P | ERTR<sub>2015</sub> | ERTR<sub>2020</sub> | P | WRTR<sub>2015</sub> | WRTR<sub>2020</sub> | P |
| Large | Yes | 33.72 | 27.63 | 0.445 | 28.30 | 44.94 | **0.018\*** | 33.72 | 28.30 | 0.445 | 27.63 | 44.94 | **0.018\*** |
|  | No | 66.28 | 72.37 |  | 71.70 | 55.06 |  | 66.28 | 71.70 |  | 72.37 | 55.06 |  |
| Medium | Yes | 53.49 | 59.21 | 0.476 | 52.83 | 48.31 | 0.572 | 53.49 | 52.83 | 1.000 | 59.21 | 48.31 | 0.156 |
|  | No | 46.51 | 40.79 |  | 47.17 | 51.69 |  | 46.51 | 47.17 |  | 40.79 | 51.69 |  |
| Small | Yes | 12.79 | 13.16 | 1.000 | 18.87 | 6.74 | **0.019\*** | 12.79 | 18.87 | 0.335 | 13.16 | 6.74 | 0.238 |
|  | No | 87.21 | 86.84 |  | 81.13 | 93.26 |  | 87.21 | 81.13 |  | 86.84 | 93.26 |  |

### NDVI values at spatio-temporal scales

NDVI comparison using one-way ANOVA showed significant differences between zones (WRTR, ERTR-core and ERTR-buffer) in both 2015 (F (2,279) = 5.81, P = 0.003) as well as 2020 (F (2,279) = 7.74, P = 0.001), at spatial scale. Subsequent post hoc test showed that WRTR and ERTR-core zones did not differ significantly in their mean NDVI values in year 2015 (P = 0.47) as well as in 2020 (P = 0.93). However, ERTR-buffer zone continuously showed significantly lower NDVI values in both the years compared to WRTR (2015: P = 0.03, 2020: P = 0.001) and ERTR-core (2015: P = 0.004, 2020: P = 0.003) zones (Figure 2c). At temporal scale, paired t-test showed that mean NDVI values improved significantly in year 2020 for WRTR (t(134) = 25.06, P < 0.0001), ERTR-core (t(66) = 9.39, P < 0.0001) as well as ERTR-buffer (t(79) = 12.45, P < 0.0001) zones compared to year 2015 (Figure 2c).

**Figure 2.**
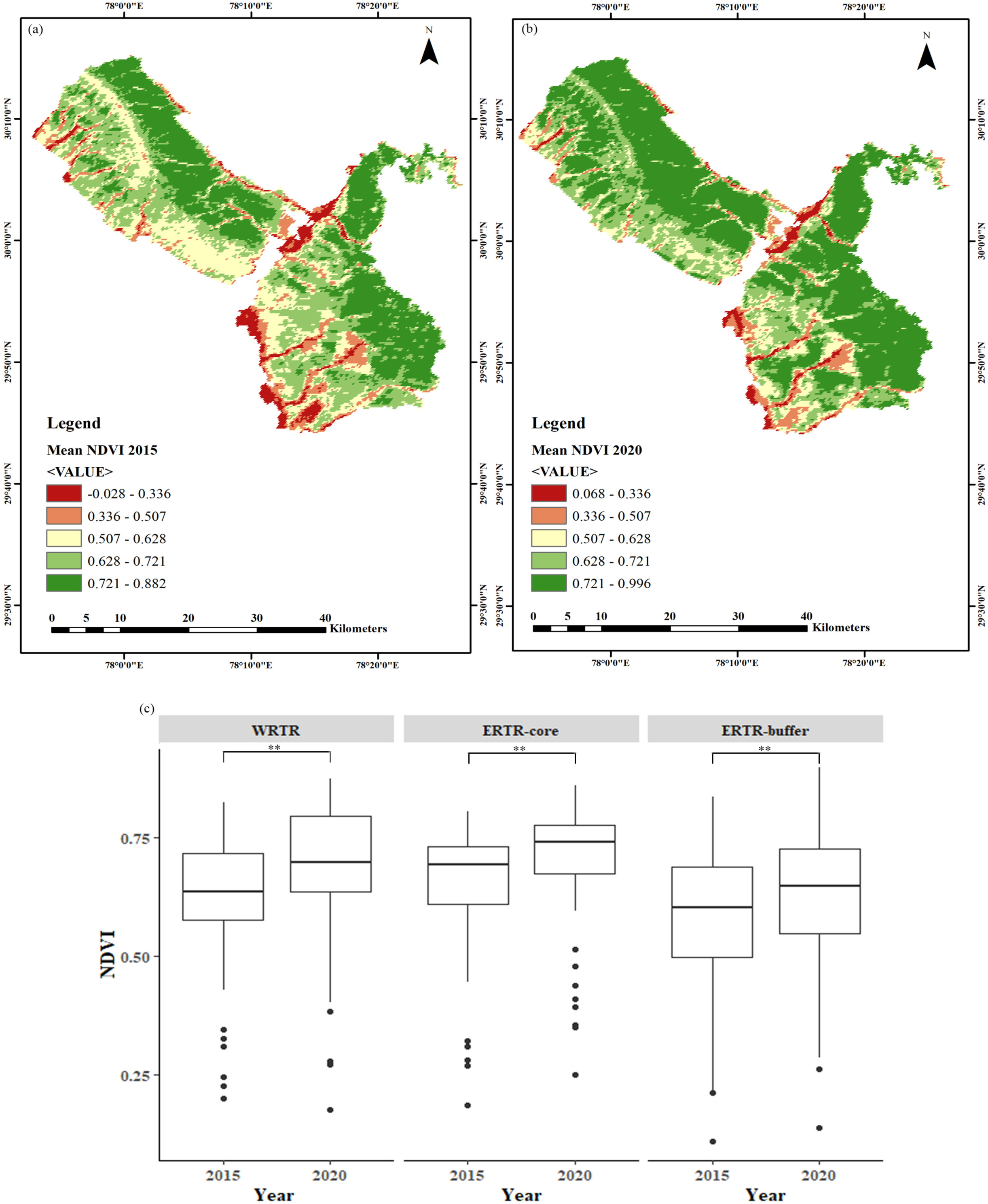
Assessment and comparison of habitat quality (through mean NDVI values) across core and buffer zones in two time points (winter 2015 and winter 2020). Panel (a) and (b) represents the mean NDVI gradients in 2105 and 2020, respectively. Panel (c) shows the temporal differences in mean NDVI values for WRTR, ERTR-core and ERTR-buffer areas. ** indicate significance value at P<0.005.

### Physiological responses of the sampled groups at spatial and temporal scales

Likelihood ratio test selected the individual GLM explanatory variables (Area*Year and Prey size, respectively) over combined model as significant factors to explain the physiological response patterns (Supplementary table 3). GLM results with Area*Year model indicated year is a significant factor (p=0.05) for fGCM data (Table 3). At spatial scale, both ERTR_2015_-WRTR_2015_ and ERTR_2020_-WRTR_2020_ comparisons showed no significant differences in mean fGCM levels (2015-z = 0.834, p = 0.838; and 2020-z = 0.253, p = 0.994) (post-hoc test, Supplementary table 2). However, temporal scale comparisons showed contrasting results where mean fGCM levels between ERTR_2015_ and ERTR_2020_ showed non-significant differences (z = −1.97, p = 0.2) but WRTR_2020_ had significantly low fGCM values than WRTR_2015_ (z = −2.73, p = 0.032) (Figure 3b(iv)). For fT3M data, GLM results showed that the area and year interaction factor (Area*Year) is a significant factor (Table 3). At spatial scale, the fT3M levels between ERTR_2015_-WRTR_2015_ showed no difference (z = −1.163, p = 0.65), but WRTR_2020_ fT3M levels were significantly higher than the ERTR_2020_ (z = 2.644, p = 0.041) (Figure 3a(iii)). At temporal scale, ERTR_2015_-ERTR_2020_ comparisons revealed no significant differences in fT3M levels (z = −1.303, p = 0.56) but WRTR_2020_ showed significantly higher values of fT3M than WRTR_2015_ (z = 2.602, p = 0.046) (Figure 3b(iii)).

**Figure 3.**
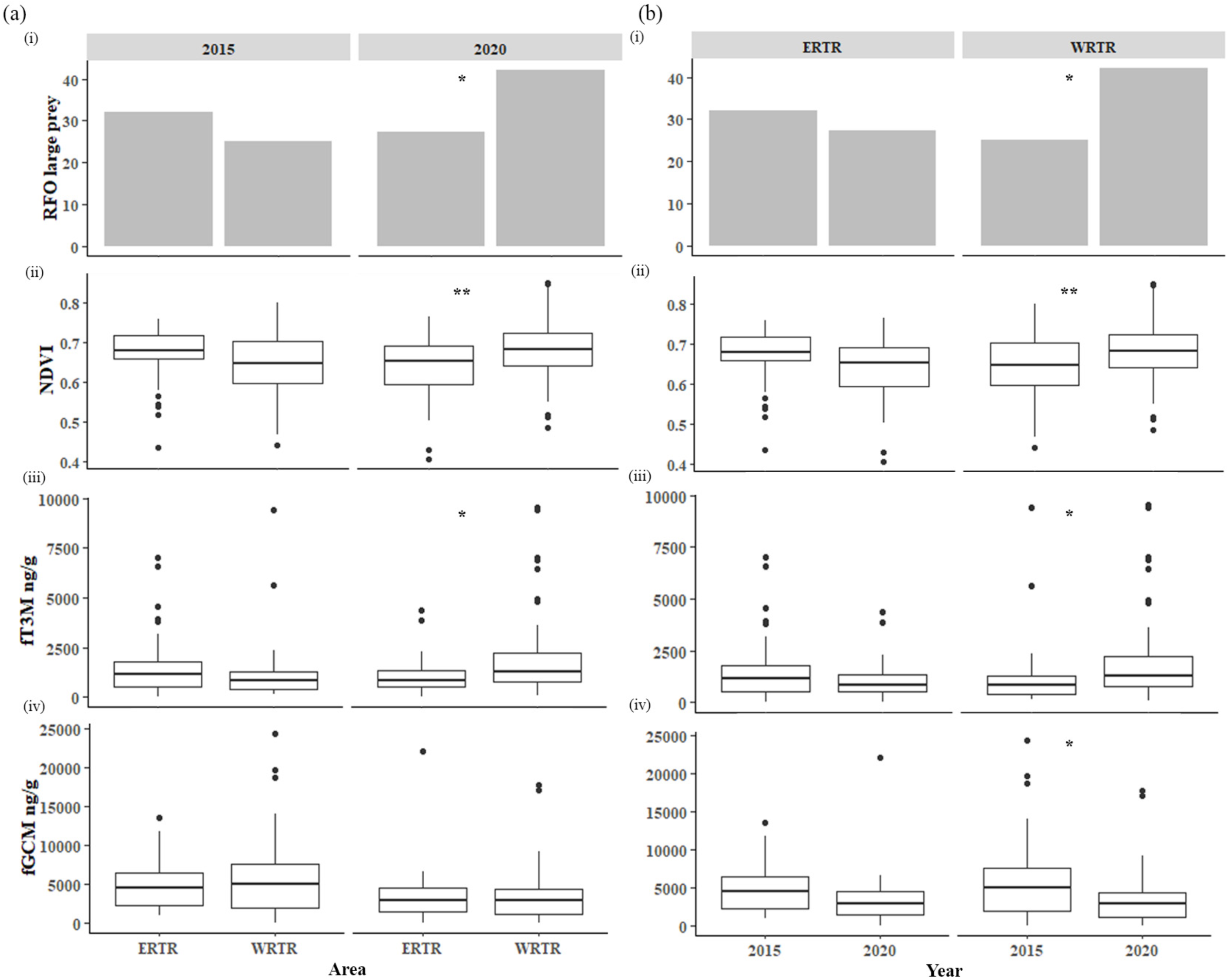
Spatio-temporal comparisons of the major variables (habitat variable-(i) large prey RFO, (ii) NDVI and physiological variable-(iii) fT3M and (iv) fGCM) used in this study. Panel (a) shows the temporal-scale differences and panel (b) indicate spatial differences. Significant differences are depicted by * (indicating P<0.05) and ** (indicating P<0.005).

**Table 3.** Results showing the association between the predictor (prey size categories and interaction of area and year) and response variables (fGCM and fT3M) based on GLM analyses. Models were fitted using log link and gamma error distributions.

| Response variable | Predictor variable | Level | Estimate | ± SE | t value | Pr(>\|t\|) |
| --- | --- | --- | --- | --- | --- | --- |
| fGCM | Area*Year | (Intercept) | 162.256 | 78.035 | 2.079 | 0.0389 * |
|  |  | AreaWest | 41.441 | 105.873 | 0.391 | 0.696 |
|  |  | Year | -0.076 | 0.039 | -1.972 | 0.0500 * |
|  |  | AreaWest:Year | -0.020 | 0.052 | -0.390 | 0.697 |
| fGCM | Prey size<br>(L, M and S) | (Intercept) | 8.377 | 0.109 | 76.946 | <2e-16 *** |
|  |  | Prey size (Medium) | -0.042 | 0.142 | -0.294 | 0.769 |
|  |  | Prey size (small) | 0.341 | 0.214 | 1.592 | 0.113 |
| fT3M | Area*Year | (Intercept) | 133.090 | 96.622 | 1.377 | 0.170 |
|  |  | AreaWest | -356.293 | 131.091 | -2.718 | 0.00718 ** |
|  |  | Year | -0.062 | 0.048 | -1.303 | 0.194 |
|  |  | AreaWest:Year | 0.177 | 0.065 | 2.719 | 0.00716 ** |
| fT3M | Prey size<br>(L, M and S) | (Intercept) | 7.538 | 0.135 | 55.876 | < 2e-16 *** |
|  |  | Prey size (Medium) | -0.458 | 0.175 | -2.612 | 0.00972 ** |
|  |  | Prey size (small) | -0.306 | 0.266 | -1.154 | 0.250 |

GLM outputs with Preysize model indicated no significant differences between prey size class (large, medium and small body-sized prey) for fGCM levels (Table 3, Figure 4a). However, the fT3M levels showed strong relationship with prey size classes, where the fT3M levels from the leopard samples with large prey remains were higher than small prey class (z =1.15, p =0.5) and significantly higher than medium prey class (z = 2.61, p = 0.02) (Table 3, Figure 4b). Linear models showed a marginally significant positive association between leopard fGCM and habitat NDVI values (t-value = 2.003, F (1, 201) = 4.01, P = 0.05) (Supplementary figure 4a), and no significant association between leopard FT3M and habitat NDVI values (t-value = −1.11, F (1, 201) = 1.23, P = 0.27) (Supplementary figure 4b).

**Figure 4.**
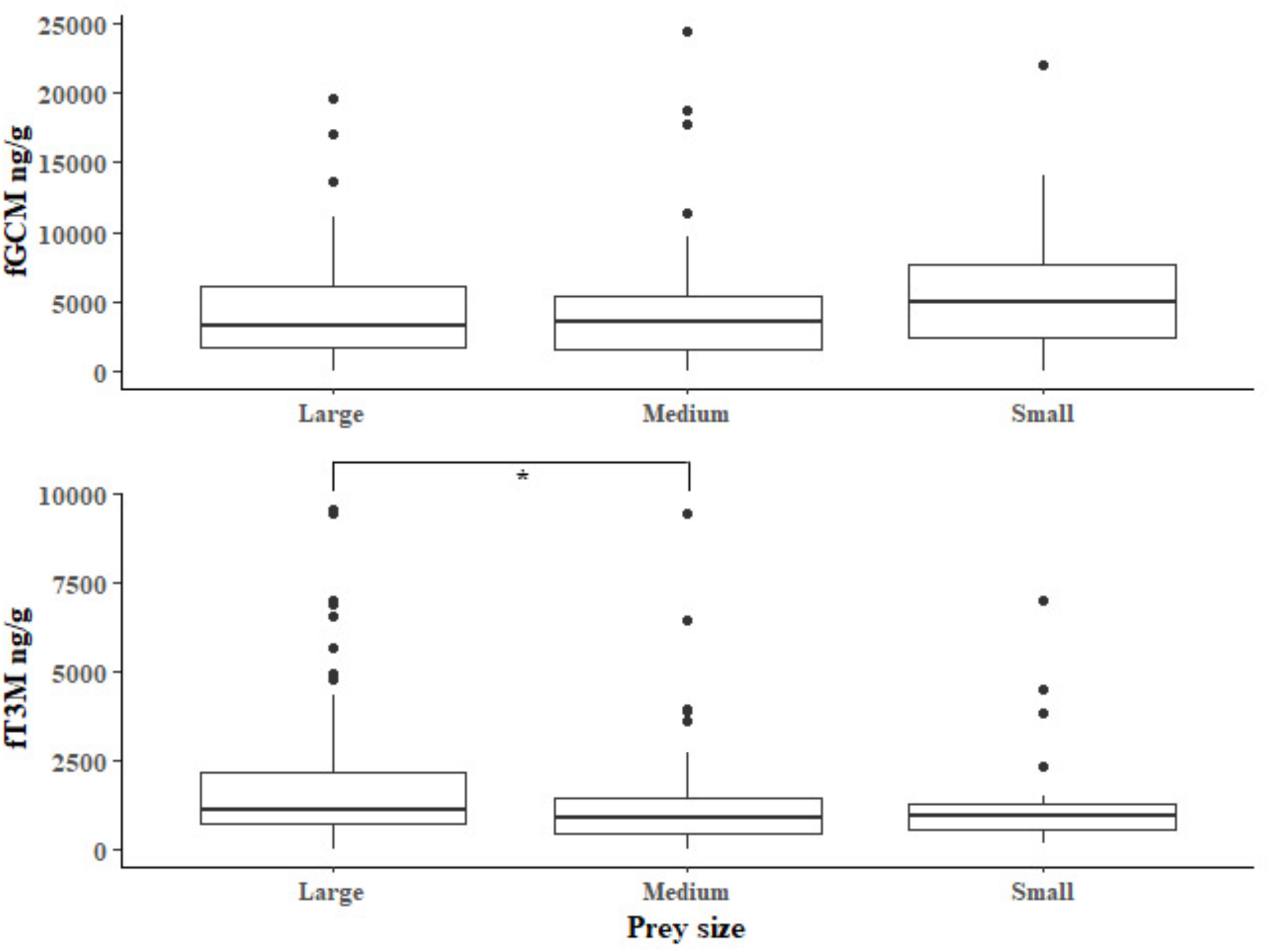
Comparison of (a) fGCM and (b) fT3M measures from faecal samples consisting large, medium and small body-sized prey. Significant differences are depicted by * (indicating P<0.05).

## 5. Discussion

As expected, our results provide strong support for certain known patterns of tiger-leopard dominance dynamics. For example, we observed one such dominance hierarchy in the form of dietary niche separation within and between ERTR and WRTR at spatio-temporal scale. In 2015, our results show relatively higher chital and similar sambar frequencies (two major prey species in the study area) in WRTR compared to ERTR (Table 1) but overall distribution of different prey size class (large, medium and small) remained similar. However, in 2020 the WRTR area showed significantly higher large-bodied and lower small-bodied prey species compared to ERTR (as hypothesized in this study). For ERTR (in 2020, spatially as well as temporally), relatively higher frequency of small-bodied prey in leopard diet can be explained through possible competitive spatial exclusion of leopards. During 2020 surveys majority of the leopard faecal samples were obtained from the buffer regions of the park, which are not prime habitats in terms of prey abundances (Harihar et al., 2020). The southern buffer and Gohri range of ERTR (see Figure 1) supports lower prey densities (25.24 individuals/ km^2^) than the core areas (39.23 individuals/ km^2^) (Harihar et al. 2020) and still hosts human settlements and livestock that exert heavy pressure on forest resources (Johnsingh et al., 1994; Harihar et al., 2014; Harihar et al., 2020) and affects the ungulates population density by hindering the forest resource availability to wild prey species (Rasal et al., 2022; Salvatori et al., 2022; Pozo et al., 2021). An alternate explanation behind such pattern could be leopard prey preferences for the medium to small bodied prey size class (Pokheral & Wegge 2019; Karanth & Sunquist 1995), but the natural experimental setup (presence of tiger in ERTR and not in WRTR) allowed us to confirm the effects of competition. The results indicate that leopard consumption of medium to small sized prey in ERTR is not driven by their preference towards these particular classes but rather is a consequence of competition from tiger. Significantly high consumption of large body-sized prey in WRTR (in absence of tiger) supports that such pattern is driven by competition pressure. Earlier Vanak et al., (2013) has also suggested similar pattern of competition (instead of preference) driven dietary niche separation

These patterns can also be explained through the results of the NDVI analyses presented here. Since the relocation of majority of the human settlements from the eastern and western RTR (from 2003 onwards) and creation of inviolate space the habitat productivity of the park has improved significantly (Harihar et al 2008). Our NDVI analyses supported this by showing significant increase in vegetation cover across the park (WRTR, ERTR-core and ERTR-buffer) (Figure 2a & b). We feel that the habitat improvement (and subsequent increase in prey density, Jhala et al., 2015; 2020) along with no-competition from tigers has resulted in availability of large-bodied prey species in the WRTR. In the ERTR, despite the habitat improvement the population faced competition pressure from tigers over years (tiger density doubled in 2020 in ERTR, Jhala et al., 2020), resulting in their potential displacement to the buffer regions (sub-optimal habitats with medium-sized prey availability). This can be substantiated by observing the faecal location patterns in this study and earlier reports of tiger and leopard occupancy in the park (Harihar et al., 2011; Rathore et al., 2015). Spatial projection of the confirmed ERTR leopard samples showed that most of the 2015 samples were distributed in the core areas whereas majority of the 2020 samples were from the buffer areas (southern boundaries of the core) (Figure 1). Given that our sampling was conducted in the same trails in the core region during both years it can be inferred that the leopard presence in ERTR core decreased in 2020, possibly due to competition from socially-dominant tigers (as seen in other Terai habitats, McDougal 1988; Wegge et al., 2009; Odden et al., 2010; Kafley et al., 2019; Thapa et al., 2021). Further, the tiger and leopard population estimation data (Jhala et al., 2020) also suggest more photographic captures of leopards at northern and southern boundaries of the ERTR-core during 2018-19. Earlier, Harihar et al (2011) and Rathore (2015) also reported a decline in leopard density in ERTR-core and their activity hotspots were concentrated towards the peripheral areas of the park as compared to the tiger activity hotspots towards the core areas. On the contrary, leopard faeces were found across more uniformly in WRTR in both sampling periods in the absence of any competition from tigers in this side of the park.

The results of the physiological impacts of dietary differences and habitat productivity largely mimicked the earlier mentioned patterns. Our measurements of fGCM and fT3M largely followed the NDVI (proxy for habitat productivity) and prey size class patterns across ERTR and WRTR, respectively (Figure 3 a & b). At spatial scale, we did not observe any significant differences in both fGCM and fT3M between ERTR and WRTR in 2015. This pattern did not support our hypothesis where we expected higher fGCM measures (from tiger competition) and lower fT3M (from possibly reduced prey accessibility) in ERTR. However, in 2020 our data show slightly different pattern in physiological measures where the fT3M levels in WRTR was significantly higher than ERTR, but no difference was found in fGCM titers. When looked at temporal scale, these patterns provide contrasting patterns for the physiological measures. Within ERTR no significant difference in fGCM and fT3M values were found between 2015 and 2020 (similar to the spatial patterns) but we observed significant differences in both measures in WRTR. The temporal-scale data provides strong support to explain the patterns in the light of ecological variables (habitat productivity and resource availability). From the observed data the influences of prey availability (associated with habitat productivity) on fT3M (nutritional hormone) can be inferred as these measures showed very similar patterns. If we consider WRTR (both 2015 and 2020) as an example, this habitat showed low NDVI values in 2015 (indicating low habitat productivity at that time due to still recovering phase after human relocation events) and corresponding lower frequency of large-body sized prey in the leopard faeces. However, during 2020 we observed drastic changes in both ecological factors where both NDVI and large prey frequencies have significantly increased. The fT3M measures exactly follow the same patterns (low in 2015 and significantly high in 2020), indicating a clear correlation between ecological variables and associated physiological responses (fT3M in this case). It is important to point out that absence of tiger in WRTR and stark differences in ecological variables in temporal scale made this inference easy compared to ERTR where more complex ecological interactions are seen. Apart from the other two ecological factors mentioned above, competition from tigers play a major role in the patterns of physiological responses in ERTR resulting in different outcomes. Here the data suggests decreasing habitat productivity and large prey frequency in leopard diet from 2015 to 2020, corroborating with a decreasing (but non-significant) trend in fT3M values between these two time periods. During 2015 the tiger density in ERTR was 2.90 (±0.87)/100 km^2^ (Jhala et al., 2015) which increased to 8 (± =1.4)/100 km^2^ (Jhala et al., 2020) during 2020, and the resulting increase in competition would explain the nutritional stress (low fT3M value) during 2020. Given the complex interactions among various ecological variables (habitat productivity, prey availability (in terms of size class) and various levels of competition pressures (from tigers)), we feel that it might take more time to observe significant differences in physiological parameters in ERTR. A similar study after 3-5 years may provide further clarification on any possible differences in fT3M values in ERTR.

One of the critical considerations in this study is the significant association of prey body size classes (large, medium and small) with fT3M measures. This was considered based on our available knowledge that large terrestrial predator distribution and abundance is significantly driven by higher biomass availability (Carbone et al., 2011) and they are known to prefer large-bodied prey due to higher energy gains (Carbone et al., 1999; 2002; 2007; 2011; Radloff & Du Toit 2004). Work on leopard energetics has also reported increased energy expenditure between meals when meal size from previous kill is large (Wilmers 2017). We used our data to test the effect of prey body size on fT3M measures, where leopard faeces with evidences of only specific body-size classes were identified and the fT3M measures were correlated (Figure 4b). In absence of any physiological validation of fT3M measures in leopard (see Mondol et al 2020 for tiger) this result can also be considered as biological validation of fT3M under field conditions.

The fGCM analyses showed slightly different patterns than the resource-driven fT3M data. Spatially, the ERTR and WRTR did not show any significant differences in fGCM levels in both 2015 and 2020 (Figure 3a(iv)) although 2020 fGCM values were relatively lower than 2015 (probably due to increased habitat productivity in 2020, at least for WRTR). However, at temporal scale the ERTR and WRTR showed contrasting patterns (in terms of our hypotheses). In ERTR, we observed a lower value of fGCM in 2020 (non-significant) when compared to 2015. This is surprising when considered that in 2020 the NDVI as well as large prey proportion was lower in ERTR (with corresponding low fT3M indicating higher nutritional stress) (Figure 3b). Physiologically this should result in high fGCM (McCormley et al., 2018; Wasser et al., 2017; 2011; Dias et al., 2017; Joly et al., 2015; Vynne et al., 2014; Ayres et al., 2012) values but the data shows an opposite pattern. While it is difficult to point out the exact mechanism behind such pattern, one possible reason could be much less competition from tigers in the buffer areas (in 2020). This could be a preferred bargain for the spatially displaced leopards from the core areas of ERTR in 2020 (tiger density 8 (±1.4)/100 km^2^ in 2020) at the cost of better nutritional status. Such behavior has been earlier documented in many anti-predatory response studies (Wasser et al., 2011; Creel et al., 2009; Ylönen et al., 2006). WRTR leopards, in absence of any competition exhibited expected physiological responses at temporal scale where we observed higher fGCM (and corresponding high nutritional stress/ low fT3M) in 2015 and low nutritional stress (high fT3M) and low fGCM in 2020. Both these patterns are also corroborating with the habitat quality and prey size class information in respective time frames (Figure 3b). It is important to point out that recent studies have cautioned regarding careful interpretations of GC/fGCM data due to contrasting pattens of directionality in GC response to chronic stress (Dickens & Romero 2013), where both increase and decrease in GC titer has been observed as a consequence of chronic stress. Therefore, this study provides strong evidence of combining additional hormones (such as T3 used here for nutritional stress) along with GC as biomarkers to reveal different physiological regulatory responses to the environment under different contexts. Multidisciplinary approaches, as used in this study would also bring out more comprehensive and ecologically meaningful outcomes that can be used in making appropriate conservation interventions.

Finally, the unique natural experimental habitat scenario, spatio-temporal sampling strategy and the patterns of fGCM and fT3M levels bring out some important conservation perspectives for leopards. Our results suggest that from a physiological perspective prey body size (large, medium and small) and availability (driven by habitat productivity) directly affect the dominant-subordinate dynamics, which is further compounded by the competition between both species (resulting in competitive exclusion). This has critical conservation implications for areas surrounding majority of the tiger landscapes across India. India has recently declared doubling its tiger numbers (population estimate of 1411 (1165-1675) in 2006 to 2967 (2603-3346) in 2018) across the country (Jhala et al. 2020). Such increase in tiger population is expected to increase pressure on sympatric leopard populations pushing them towards buffers or more sub-optimal habitats, further exacerbating chances of human-leopard conflict. Therefore, expansion of leopard monitoring and population estimation efforts to buffers, their management in the context of conflicts and understanding of local factors driving the changes in population pattern would be critical for their future conservation. Our results highlight the importance of good-quality habitats and prey base for this species and future conservation efforts should ensure availability of the same for their persistence. This study also emphasizes the importance of similar work on other carnivore guilds particularly in the context of the ongoing Anthropocene, which is affecting inter-species dynamics globally.

## Acknowledgement

We acknowledge the Uttarakhand Forest Department for providing necessary permits to carry out the research. Our thanks to the Forest Department officials and frontline staff members for their support and assistance during field surveys. We acknowledge help from Shrutarshi, Tista, Shrushti, Harshvardhan, Ramesh, Satish, Annu, Bhura, Ranjhu, Abbi, Ammi, Inam, Imam and Anubhuti for their help during sampling, Bhoomika for assisting in laboratory work and Meghna for analytical help. We thank the Director, Dean, Research Coordinator and TRAC members of Wildlife Institute of India for their constant support during implementation this work.

## Funding

This research was funded by Grant-in-Aid support from Wildlife Institute of India to Samrat Mondol. Samrat Mondol was supported by Department of Science and Technology INSPIRE Faculty Award (IFA12-LSBM-47).

## Conflict of interests

The authors declare no conflict of interests.

## Author contributions

**Shiv Kumari Patel:** Conceptualization, Data generation, Data curation, Formal analysis, Validation, Visualization, Writing-Original Draft, Writing-Review and Editing; **Sourabh Ruhela:** Data generation, Data curation, Writing-Review and Editing; **Suvankar Biswas:** Data generation, Data curation, Analytical discussions, Writing-Review and Editing; **Supriya Bhatt:** Data generation, Writing-Review and Editing; **Bivash Pandav:** Conceptualization, Writing-Review and Editing, Supervision. **Samrat Mondol:** Conceptualization, Data Curation, Analysis, Resources, Writing-Original draft, Writing-Review and Editing, Supervision, Project administration, Funding acquisition.

---

**Supplementary figure 1.**
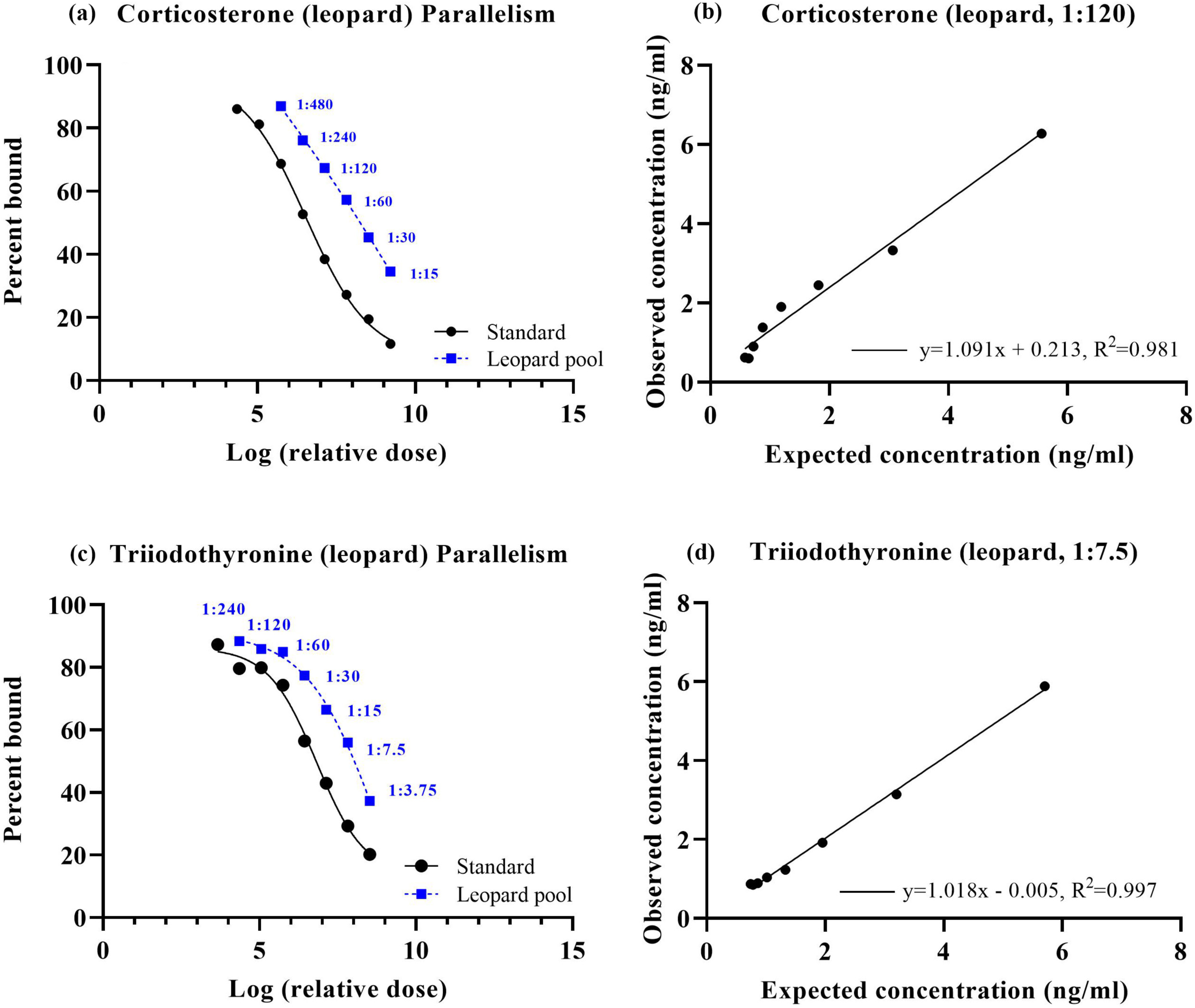
Parallelism and accuracy plots for faecal corticosterone and T3 EIA assays.

**Supplementary figure 2.**
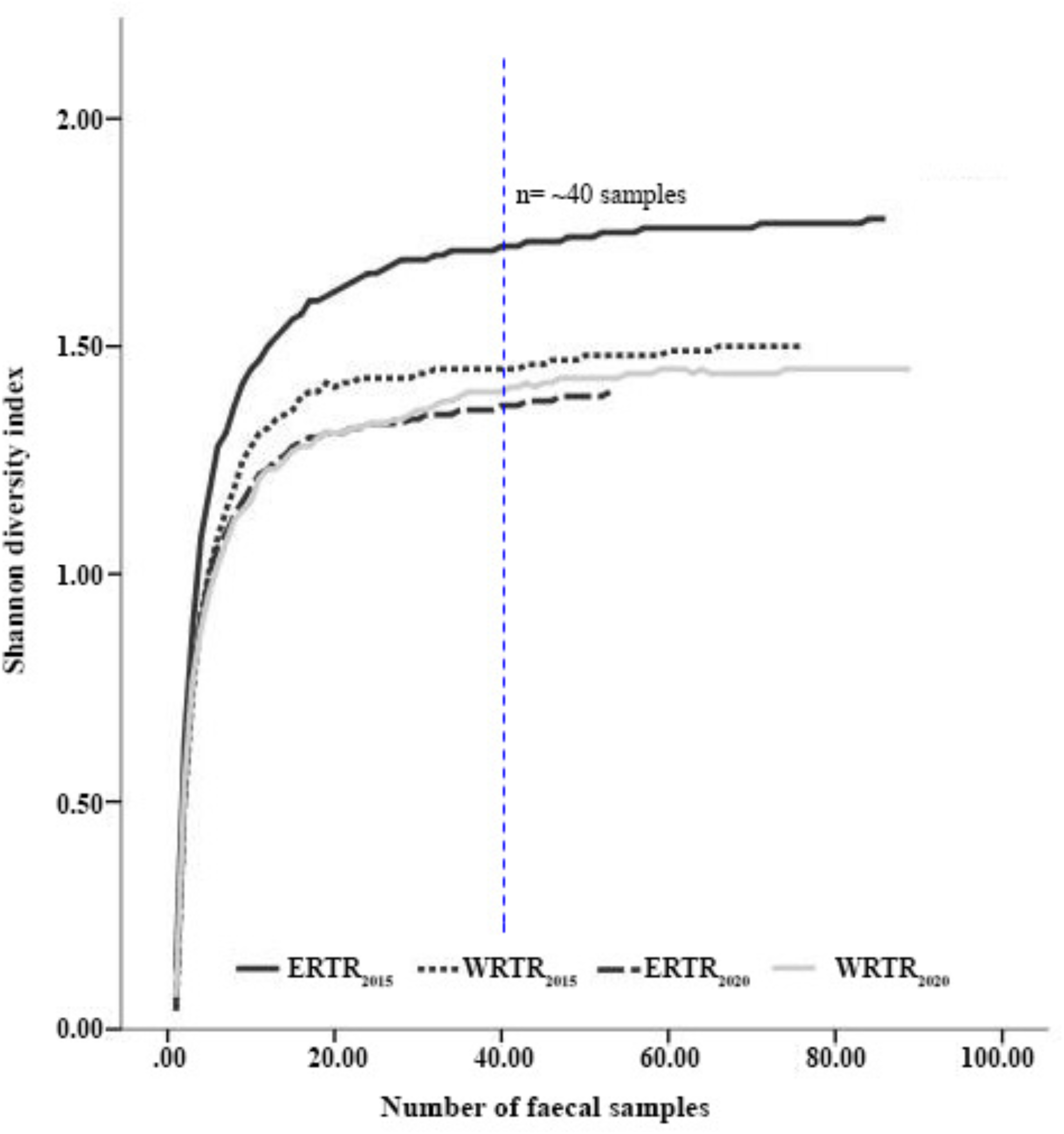
The rarefaction curves indicating sample sufficiency for leopard food habit assessments. Different line types represent separate spatial (ERTR and WRTR) and temporal (2015 and 2020) sample combinations. The curves stabilize at a sample size of ∼40, represented by dashed blue line.

**Supplementary figure 3.**
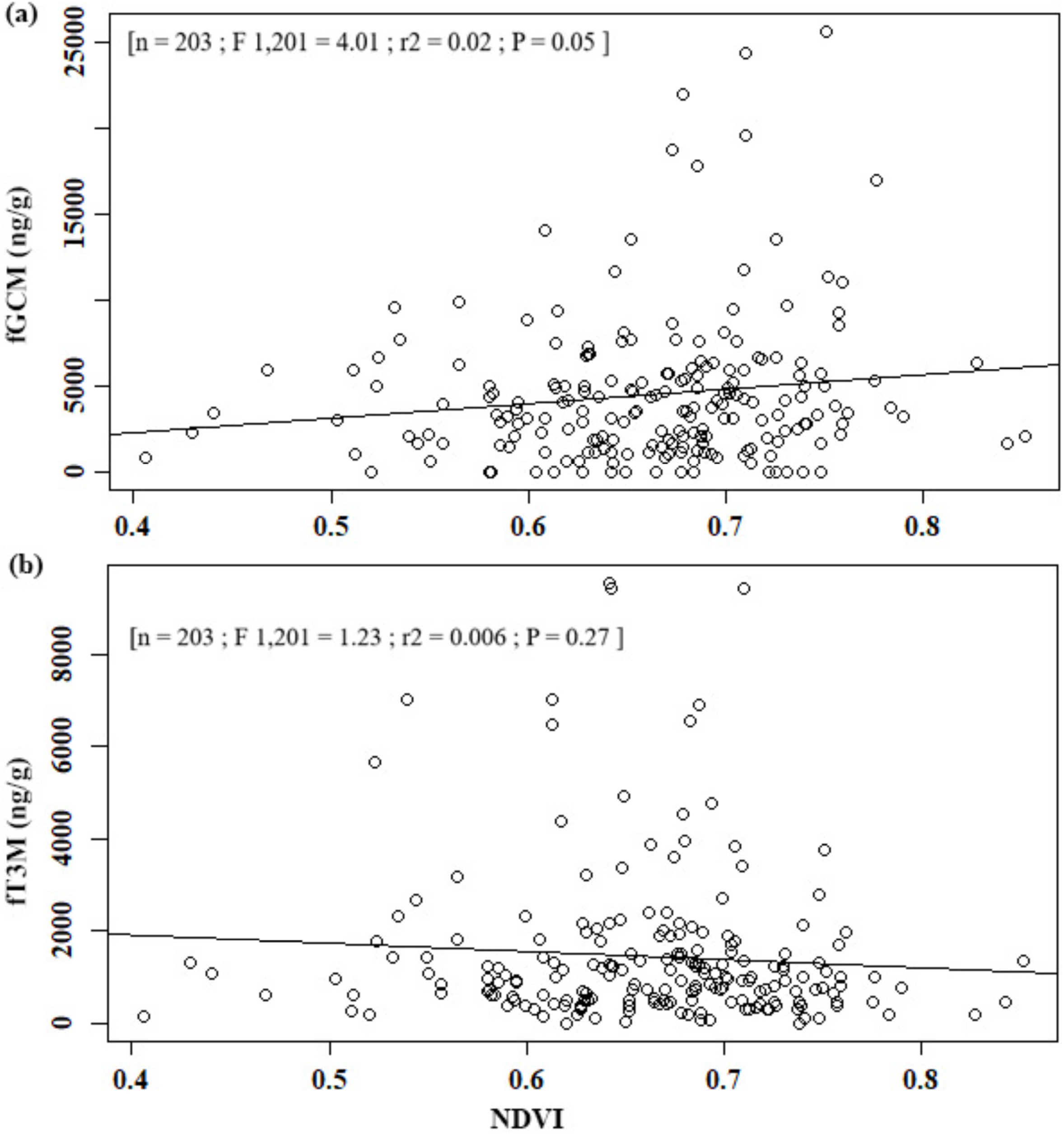
Relationship of habitat NDVI values with leopard fGCM and fT3M levels. Scatter-plots showing (a) the positive association between NDVI values and fGCM levels and b) no significant association between NDVI values and fT3M levels.

**Supplementary Table 1:**
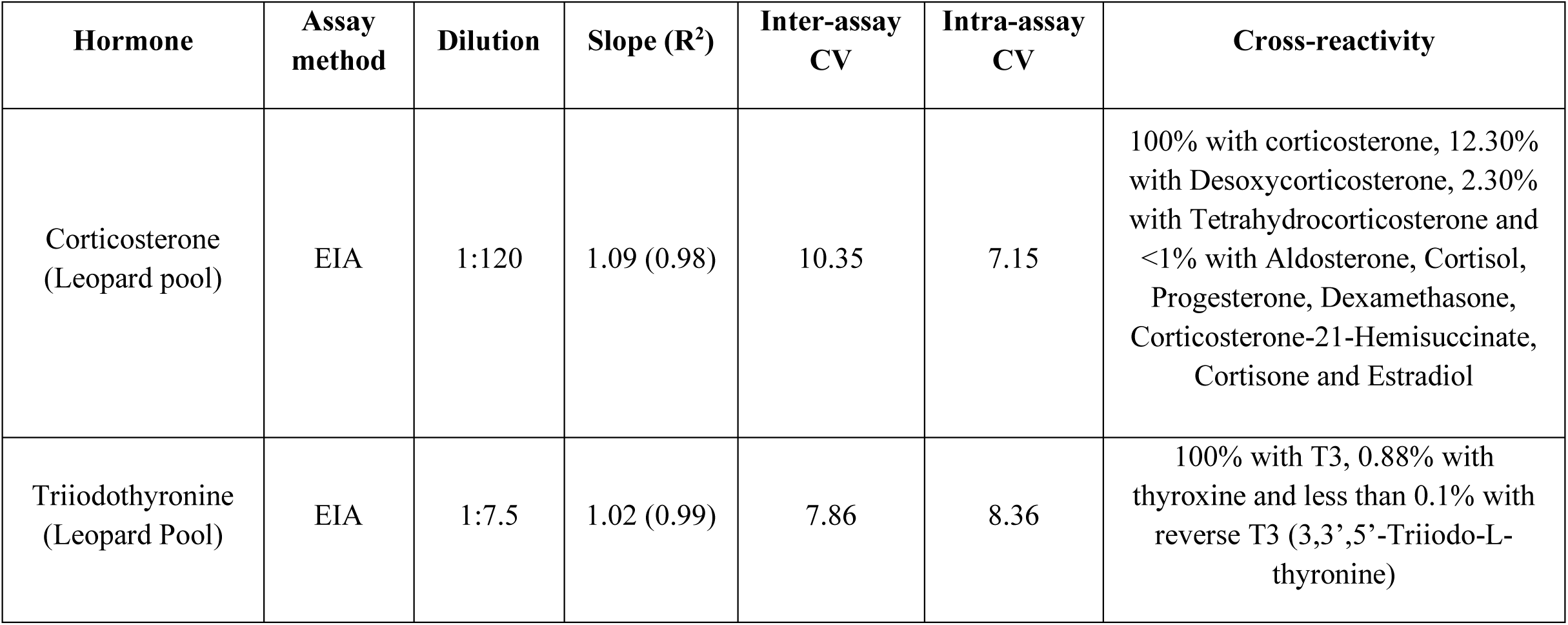
Details of the faecal hormone assays conducted for leopards in this study.

| Hormone | Assay method | Dilution | Slope (R<sup>2</sup>) | Inter-assay CV | Intra-assay CV | Cross-reactivity |
| --- | --- | --- | --- | --- | --- | --- |
| Corticosterone (Leopard pool) | EIA | 1:120 | 1.09 (0.98) | 10.35 | 7.15 | 100% with corticosterone, 12.30% with Desoxycorticosterone, 2.30% with Tetrahydrocorticosterone and <1% with Aldosterone, Cortisol, Progesterone, Dexamethasone, Corticosterone-21-Hemisuccinate, Cortisone and Estradiol |
| Triiodothyronine (Leopard Pool) | EIA | 1:7.5 | 1.02 (0.99) | 7.86 | 8.36 | 100% with T3, 0.88% with thyroxine and less than 0.1% with reverse T3 (3,3',5'-Triiodo-L-thyronine) |

**Supplementary Table 2.** Results of Post hoc pair wise comparisons.

|  | **Post hoc (Tukey HSD)** | **Estimate** | **Std. Error** | **z value** | **Pr(>\|z\|)** |
| --- | --- | --- | --- | --- | --- |
| fGCM (Area*Year) | ERTR<sub>2020</sub> - ERTR<sub>2015</sub> | -0.382 | 0.193 | -1.972 | 0.198 |
|  | WRTR<sub>2020</sub> - WRTR<sub>2015</sub> | -0.484 | 0.177 | -2.729 | 0.0317 * |
|  | WRTR<sub>2015</sub> - ERTR<sub>2015</sub> | 0.151 | 0.181 | 0.834 | 0.838 |
|  | WRTR<sub>2020</sub> - ERTR<sub>2020</sub> | 0.048 | 0.190 | 0.253 | 0.994 |
| fGCM (Prey size) | Medium - Large | -0.042 | 0.142 | -0.294 | 0.952 |
|  | Small - Large | 0.341 | 0.214 | 1.592 | 0.243 |
|  | Small - Medium | 0.383 | 0.206 | 1.863 | 0.146 |
| fT3M (Area*Year) | ERTR<sub>2020</sub> - ERTR<sub>2015</sub> | -0.312 | 0.240 | -1.303 | 0.560 |
|  | WRTR<sub>2020</sub> - WRTR<sub>2015</sub> | 0.571 | 0.220 | 2.602 | 0.0456 * |
|  | WRTR<sub>2015</sub> - ERTR<sub>2015</sub> | -0.260 | 0.224 | -1.163 | 0.649 |
|  | WRTR<sub>2020</sub> - ERTR<sub>2020</sub> | 0.623 | 0.236 | 2.644 | 0.0408 * |
| fT3M (Prey size) | Medium - Large | -0.458 | 0.175 | -2.612 | 0.024 * |
|  | Small - Large | -0.306 | 0.266 | -1.154 | 0.475 |
|  | Small - Medium | 0.152 | 0.255 | 0.596 | 0.819 |

**Supplementary Table 3.** Results of Likelihood ratio test (LRT).

| Likelihood ratio test |  |
| --- | --- |
| Models | Pr(>Chi) |
| fGCM ~ Prey size | 0.006** |
| fGCM ~ Prey size + Area * Year |  |
| fGCM ~ Area*Year | 0.117 |
| fGCM ~ Prey size + Area * Year |  |
| fT3M ~ Prey size | 0.092 |
| fT3M ~ Prey size + Area * Year |  |
| fT3M ~ Area*Year | 0.164 |
| fT3M ~ Prey size + Area * Year |  |

**Supplementary Table 4.** Influence of NDVI values on fGCM and fT3M levels in leopards based on linear models.

| Response | Predictor | Estimate | ±SE | t value | Pr (>\|t\|) |
| --- | --- | --- | --- | --- | --- |
| fGCM | (Intercept) | -1014 | 2764 | -0.367 | 0.714 |
|  | NDVI | 8309 | 4149 | 2.003 | 0.0465* |
| fT3M | (Intercept) | 2630 | 1075 | 2.447 | 0.0153 |
|  | NDVI | -1786 | 1613 | -1.107 | 0.2697 |

lm (formula = fGCM~ NDVI, data); lm (formula = fT3M~ NDVI, data).

**Supplementary Table 5.** Results showing the output of two-way ANOVA, showing prey RFO and biomass comparison within habitat across all compared groups.

| S.No. | variable | Groups compared | F statistic | P-value |
| --- | --- | --- | --- | --- |
| 1 | Prey RFO | ERTR<sub>2015</sub> vs. WRTR<sub>2015</sub> | 40.39 | 0.00001 |
| 2 | Prey RFO | ERTR<sub>2020</sub> vs. WRTR<sub>2020</sub> | 21.49 | 0.0001 |
| 3 | Prey RFO | ERTR<sub>2015</sub> vs. ERTR<sub>2020</sub> | 31.16 | 0.00003 |
| 4 | Prey RFO | WRTR<sub>2015</sub> vs. WRTR<sub>2020</sub> | 30.74 | 0.00003 |
| 5 | Prey biomass | ERTR<sub>2015</sub> vs. WRTR<sub>2015</sub> | 19.49 | 0.0001 |
| 6 | Prey biomass | ERTR<sub>2020</sub> vs. WRTR<sub>2020</sub> | 24.61 | 0.00007 |
| 7 | Prey biomass | ERTR<sub>2015</sub> vs. ERTR<sub>2020</sub> | 29.92 | 0.00003 |
| 8 | Prey biomass | WRTR<sub>2015</sub> vs. WRTR<sub>2020</sub> | 27.92 | 0.00004 |

## Notes

### Competing Interest Statement

The authors have declared no competing interest.

